# Akaluc bioluminescence offers superior sensitivity to track *in vivo* dynamics of SARS-CoV-2 infection

**DOI:** 10.1101/2023.10.12.561993

**Authors:** Tomokazu Tamura, Hayato Ito, Shiho Torii, Lei Wang, Rigel Suzuki, Shuhei Tusjino, Akifumi Kamiyama, Yoshitaka Oda, Yuhei Morioka, Saori Suzuki, Kotaro Shirakawa, Kei Sato, Kumiko Yoshimatsu, Yoshiharu Matsuura, Satoshi Iwano, Shinya Tanaka, Takasuke Fukuhara

## Abstract

Monitoring *in vivo* viral dynamics can improve our understanding of pathogenicity and tissue tropism. For positive-sense, single-stranded RNA viruses, several studies have attempted to monitor viral kinetics *in vivo* using reporter genomes. The application of such recombinant viruses can be limited by challenges in accommodating bioluminescent reporter genes in the viral genome. Conventional luminescence also exhibits relatively low tissue permeability and thus less sensitivity for visualization *in vivo*. Here we show that unlike NanoLuc bioluminescence, the improved method, termed AkaBLI, allows visualization of severe acute respiratory syndrome coronavirus 2 (SARS-CoV-2) infection in Syrian hamsters. By successfully incorporating a codon-optimized Akaluc luciferase gene into the SARS-CoV-2 genome, we visualized *in vivo* infection, including the tissue-specific differences associated with particular variants. Additionally, we could evaluate the efficacy of neutralizing antibodies and mRNA vaccination by monitoring changes in Akaluc signals. Overall, AkaBLI is an effective technology for monitoring viral dynamics in live animals.

## Introduction

Since the discovery of firefly luciferase (FLuc), bioluminescence has become a powerful tool^1^ for *in vivo* studies of gene regulation, particularly bioluminescence imaging (BLI), which enables spatiotemporal monitoring of live animals. BLI has been utilized to monitor transgene expression, tumor growth, metastasis, and progression of infection.^2^ In virology, BLI can be harnessed for investigating pathogenesis, immune responses to infection, and the efficacy of therapies *in vivo*. Indeed, several studies have used recombinant viruses containing bioluminescent reporter genes to determine viral kinetics (reviewed in ^3^). However, for positive-sense RNA viruses, creating such reporter strains can be challenging due to the limited loci within the viral genome that can accommodate these insertions. In addition, conventional luminescence exhibits relatively low tissue permeability. Thus, when studying infection *in vivo*, viral replication in the deep organs, such as the brains and lungs, can be difficult to visualize. FLuc and D-luciferin was the first enzyme-substrate combination for which bioluminescent signals could be measured in mice.^4^ Of the approximately 30 naturally occurring bioluminescence systems that have been discovered,^5^ FLuc and nanoluciferase (NanoLuc) are most commonly used for BLI. Compared to FLuc, NanoLuc offers a 150-fold increase in luminescence, is smaller, and has enhanced stability.^6^ However, the short emission wavelength of NanoLuc makes it difficult to measure signal *in vivo* from deep organs such as the lungs. Similarly, FLuc emission also has weak tissue permeability.

To overcome this, Akaluciferase (Akaluc) and its substrate AkaLumine, derived from FLuc and D-luciferin, respectively, produce approximately 52 times more detectable light per reporter molecule from intrapulmonary locations.^7^ Using AkaBLI, individual tumor cells expressing Akaluc can be visualized in live animals. Here, we engineered a recombinant severe acute respiratory syndrome coronavirus-2 (SARS-CoV-2) carrying the Akaluc gene and investigated its *in vivo* utility. SARS-CoV-2 is a single-stranded, positive-sense RNA virus of the *Coronaviridae* family and the causative agent of COVID-19.^8^ As SARS-CoV-2 has circulated in human populations, new variants have repeatedly emerged,^9^ sparking continued research. In a previous collaborative study, we found evidence that a SARS-CoV-2 variant of the Omicron lineage had evolved to have altered usage of the serine protease TMPRSS2,^10^ exhibiting a different pattern of replication.

In the present study, we generated a recombinant SARS-CoV-2 encoding Akaluc and the spike (S) protein of the early pandemic lineages B.1.1 and B.1.351 (Beta); the Omicron subvariant BA.1; or the currently circulating Omicron subvariant XBB.1.5. Then, we investigated whether AkaBLI would reflect the differences in tissue tropism that have been observed for select variants. In addition, we used AkaBLI to evaluate the protection conferred by licensed neutralizing monoclonal antibodies^11^ and mRNA vaccine^12^ against SARS-CoV-2.

## Results

### Characterization of SARS-CoV-2 carrying the Akaluc luciferase gene *in vitro*

First, we produced cDNA clones of SARS-CoV-2 B.1.1 containing the wild-type Akaluc gene in place of viral open reading frames (ORFs) 6-8, which encode accessory proteins (**Figure 1A**).^13^ However, infectious virus could not be recovered from transfected cells (**Figure 1B**). As the GC content of SARS-CoV-2 is relatively low (ca. 38 %), we generated a mutant Akaluc carrying synonymous substitutions to optimize the GC content (**Figure S1**). As shown in **Figure 1B**, cytopathic effects were observed in the cells transfected with SARS-CoV-2 containing this optimized Akaluc, suggesting that the genetic fitness of foreign genes is critical for viral propagation.

**Figure 1.**
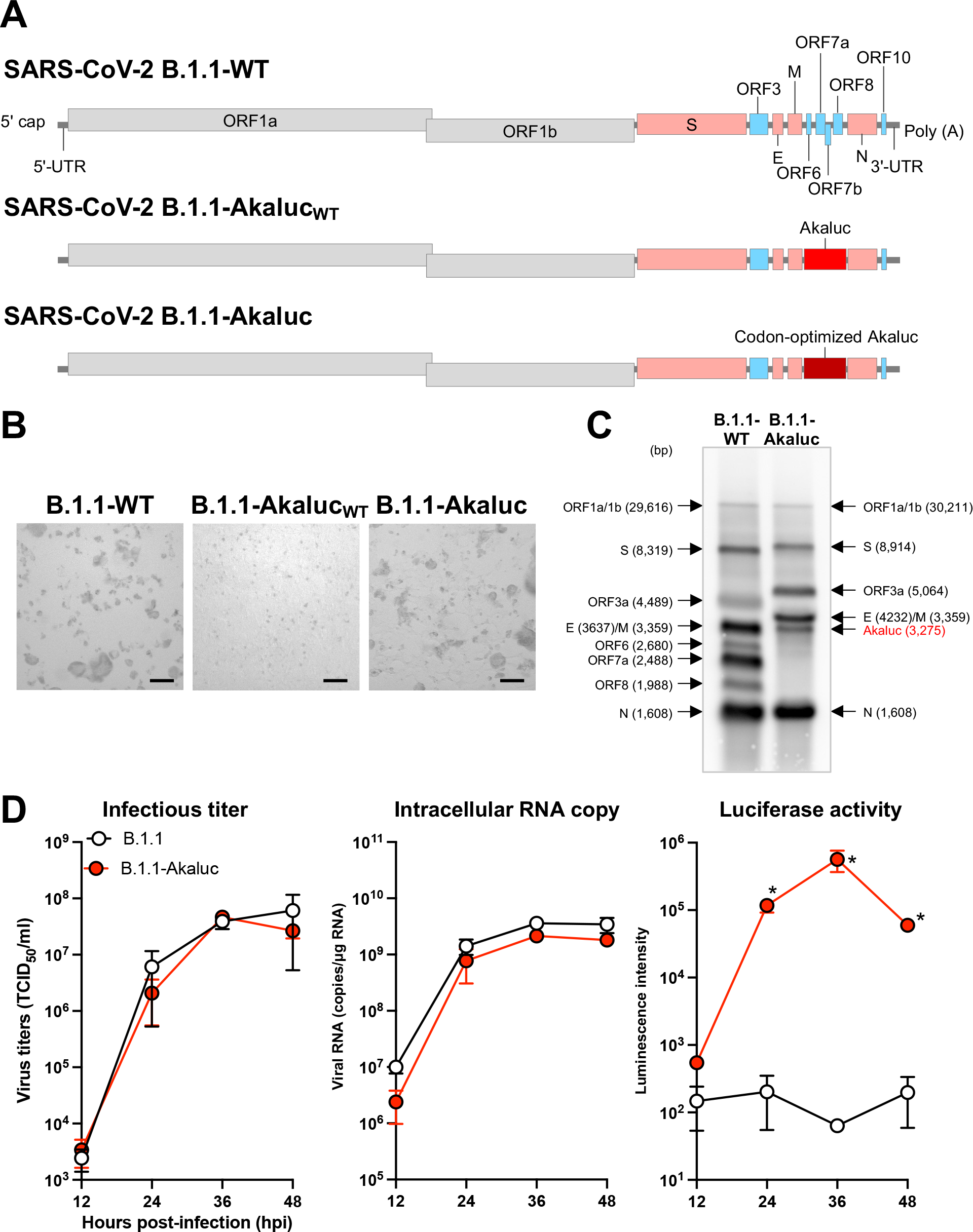
The virological features of SARS-CoV-2 Akaluc *in vitro*. **(A)** The gene structures of wild-type SARS-CoV-2 (B.1.1) and SARS-CoV-2 carrying either the wildtype or codon-optimized Akaluc luciferase gene in place of ORFs 6-8 (B.1.1-Akaluc_WT_ or B.1.1-Akaluc, respectively). **(B)** Viral production was judged by observation of cytopathic effects upon transfection with the transfection of the 3 respective genomes. Representative images are shown from 6 days post transfection. Scale bars:100 µm. **(C)** Northern blot analysis of subgenomic RNAs (sgRNAs). RNA was extracted from VeroE6/TMPRSS2 cells infected with B.1.1 or B.1.1-Akaluc and subjected to northern blot analysis. Black arrows indicate the bands for each sgRNA with size in parentheses. **(D)** Growth kinetics of B.1.1 and B.1.1-Akaluc *in vitro*. VeroE6/TMPRSS2 cells were infected with either B.1.1 or B.1.1-Akaluc (MOI = 0.001). Infectious titers in the culture supernatants (left panel), the copy number of intracellular viral RNA (middle panel), and the luciferase activity (right panel) were determined at the indicated timepoints. Asterisks indicate significant differences (*, *p* < 0.05) with the results of the wildtype virus. Assays were performed independently in duplicate.

The resultant virus, termed B.1.1-Akaluc, was subjected to northern blot analyses to confirm viral RNA synthesis. As expected, since ORFs 6-8 were replaced with the codon-optimized Akaluc, only five, instead of eight, subgenomic RNAs were detected in cells infected with B.1.1-Akaluc. The four subgenomic RNAs from ORF1a/1b and S were larger than those in the parental strain, one RNA from Akaluc gene and the other RNAs were similar in size (**Figure 1C**), indicating the Akaluc gene was incorporated into the viral genome and successfully maintained in descendent viral RNA. To investigate the viral properties of B.1.1-Akaluc in cell culture, VeroE6 cells expressing human TMPRSS2 (VeroE6/TMPRSS2 cells^14^) were infected with either B.1.1 or B.1.1-Akaluc at a multiplicity of infection (MOI) of 0.001, and the replication kinetics of the viruses were evaluated. The infectious titers of culture supernatants (**Figure 1D**, left panel) and intracellular viral RNA levels (**Figure 1D**, middle panel) were comparable over time for the two viruses. As expected, luciferase activity (**Figure 1D**, right panel) was only detected in cultures infected with B.1.1-Akaluc, mirroring the kinetics observed for intracellular viral RNA. These results indicated that replacing ORFs 6-8 of the B.1.1 genome with the Akaluc gene did not affect viral replication or production of infectious particles.

Next, to examine the stability of the reporter gene in B.1.1-Akaluc, we serially passaged the virus in HEK293-3P6C33 cells (express ACE2 and TMPRSS2)^15^ for five rounds. As shown by PCR amplification using virus-specific primers and by sequencing (**Figure S2A**), the Akaluc gene was steadily maintained, suggesting that the codon-optimized Akaluc gene was stable in the B.1.1 genome.

### Evaluation of *in vivo* viral dynamics of SARS-CoV-2 by AkaBLI

Syrian hamsters support SARS-CoV-2 infection and are thus used as a small animal model for *in vivo* study.^16^ First, to confirm the pathogenicity of the Akaluc recombinant virus *in vivo*, we inoculated hamsters intranasally with either B.1.1 or B.1.1-Akaluc (*n*=6 per group). Weight was monitored daily for 7 days post-infection (dpi) (**Figure 2A**).^17-24^ Consistent with our previous studies,^17,20^ uninfected hamsters gained weight daily. In contrast, the infected hamsters exhibited weight loss, with the decline in weight greater in hamsters infected with B.1.1 vs those infected with B.1.1-Akaluc. Viral RNA could be detected in oral swabs collected from infected hamsters, but the copy number was higher with B.1.1 infection compared to B.1.1-Akaluc (**Figure S2B**). These observations suggest that the B.1.1-Akaluc was still able to replicate in hamsters but with slightly attenuated pathogenicity, likely due to insertion of the foreign gene.

**Figure 2.**
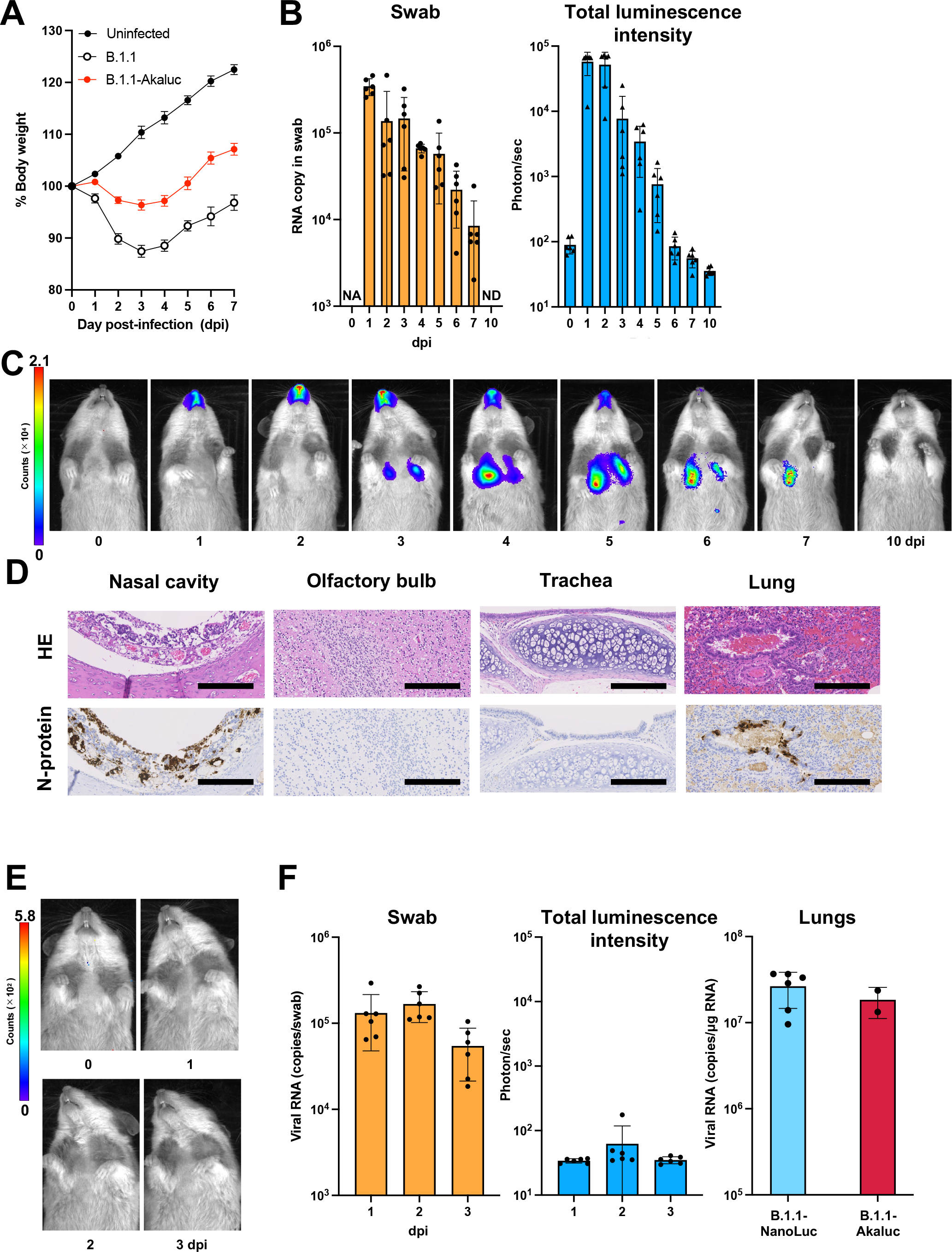
The virological features of B.1.1-Akaluc *in vivo*. **(A)** Syrian hamsters (*n*=6 per group) were intranasally inoculated with saline, B.1.1, or B.1.1-Akaluc. Body weight was measured daily through 7 days post-infection (dpi). **(B)** Syrian hamsters (*n*=6) were intranasally inoculated with B.1.1-Akaluc. Viral RNA copies in the oral swab (left panel) and the luminescence intensity of the nasal cavity of hamsters (right panel) inoculated with B.1.1-Akaluc were measured every 24 h through 7 dpi and then again at 10 dpi. **(C)** *In vivo* bioluminescence imaging of hamsters inoculated with B.1.1-Akaluc was performed daily through 7 dpi and again at 10 dpi. A representative image from an infected hamster is shown. **(D)** IHC of the viral N protein (stained brown) in the nasal cavity, olfactory bulb, trachea, and lungs of hamsters at 3 dpi with B.1.1-Akaluc. Representative figures are shown. Scale bars: 200 µm. HE, hematoxylin and eosin (**E, F**) Syrian hamsters (*n*=6) were inoculated with B.1.1 carrying the NanoLuc luciferase gene (B.1.1-NanoLuc). Bioluminescence imaging of hamsters was performed daily through 3 dpi and a representative image from an infected hamster is shown (**E**). Oral swabs were collected (**F**, left panel) and the luminescence intensity of the nasal cavity (**F**, middle panel) measured daily through 3 dpi. The lung hilum was harvested at 3 dpi and for quantification of viral RNA (**F**, right panel). Two hamsters infected with B.1.1-Akaluc served as controls.

Next, to investigate the utility of B.1.1-Akaluc for *in vivo imaging*, we assessed Akaluc signals every 24 h up to 7 dpi and again at 10 dpi by IVIS Spectrum Imaging (**Figures 2B** and **2C**). At 1 dpi, signal was detected in the nasal cavity and then in the lungs from days 3 to 7 post-infection (**Figure 2C**), indicating that B.1.1-Akaluc was able to spread through the trachea and reach the lungs after intranasal injection. Starting at 6 dpi, signal was no longer observed in the nasal cavity and by 10 dpi, not in the lungs either. Importantly, the viral RNA copies in oral swabs corresponded with the strength of the Akaluc signals measured in the nasal cavity (**Figure 2B**).

To further investigate the relationship between the Akaluc signals and viral replication in hamsters, viral nucleocapsid (N) protein was assessed by immunohistochemical (IHC) analysis in formalin-fixed organs, including the nasal cavity, olfactory bulb, trachea, and lungs, at 3 dpi. In the hamsters infected with B.1.1-Akaluc, N-positive cells were observed in the nasal cavity and the alveolar space around the bronchi/bronchioles of the lungs (**Figure 2D**). In contrast, N-positive cells were rarely detected in the olfactory bulb and epithelial cells of the trachea. These histological findings reflected the location of the signals visualized by AkaBLI.

NanoLuc has been used for *in vivo* imaging in studies of SARS-CoV-2 in transgenic mice expressing human ACE2 (hACE2)^25,26^ and humanized mice engrafted with human fetal lung xenografts in the skin.^27^ To assess whether NanoLuc signals can be used in hamsters, we generated recombinant B.1.1 carrying the NanoLuc gene (B.1.1-NanoLuc) in the same cassette as previously reported.^28^ Although B.1.1-NanoLuc replicated efficiently, with strong, specific signals observed in cell culture using IVIS Spectrum imaging (**Figure S2C**), no NanoLuc signals were detected in hamsters when using the fluorofurimazine (FFz) substrate which is developed for *in vivo* usage^29^ (**Figure 2E**) despite viral RNA being serially detected in oral swabs (**Figure 2F**). When we euthanized the inoculated animals and then directly administered FFz into the pleural cavity, the NanoLuc signals were able to be visualized (**Figure S2D**). Viral RNA copies in the lungs of the B.1.1-NanoLuc–infected hamsters were comparable to those in the B.1.1-Akaluc–infected hamsters at 3 dpi. Overall, these data indicate that Akaluc, but not NanoLuc, allows for visualization of the *in vivo* dynamics of SARS-CoV-2 infection in the respiratory system, including the lungs, of hamsters.

As shown in numerous studies, SARS-CoV-2 evolves as it circulates among humans.^8^ The Omicron variants have altered usage of TMPRSS2, the receptor for viral entry, and consequently replicate more in the upper respiratory system compared to ancestral SARS-CoV-2.^10^ To investigate whether AkaBLI can assess the differences in infection of SARS-CoV-2 variants, we generated two additional recombinant viruses carrying the S gene from the ancestral lineage B.1.351.1 and the Omicron lineage BA.1, (B.1.351.1-Akaluc and BA.1-Akaluc, respectively). As observed for B.1.1-Akaluc, B.1.351.1-Akaluc and BA.1-Akaluc replicated and exhibited Akaluc signals in cell culture infections (**Figure S2E**). As shown in **Figure 3A**, the two Omicron variants carrying Akaluc replicated in hamsters, with Akaluc signal detectable starting at 1 dpi. B.1.351.1-Akaluc displayed similar distribution of Akaluc signal as B.1.1-Akaluc, with signals detected in the lungs at 3 dpi. In contrast, BA.1-Akaluc signals remained limited to the nasal cavity at 3 dpi. The strength of Akaluc signals corresponded to viral RNA copies in the oral swabs (**Figure 3B**). Consistent with the BLI observations, the viral RNA copies in the lungs of BA.1-Akaluc–infected hamsters were significantly lower than those of the B.1.351.1-Akaluc–infected hamsters (**Figure 3C**). N-positive cells as detected by IHC also corresponded with the Akaluc signals observed in the different tissues analyzed. As with the Akaluc signal, few N-positive cells were observed in the lung section of BA.1-Akaluc–infected hamsters compared with that of B.1.351.1-Akaluc–infected hamsters (**Figure 3D**). These data indicate that the Akaluc signals can reflect the replication properties of SARS-CoV-2 variants.

**Figure 3.**
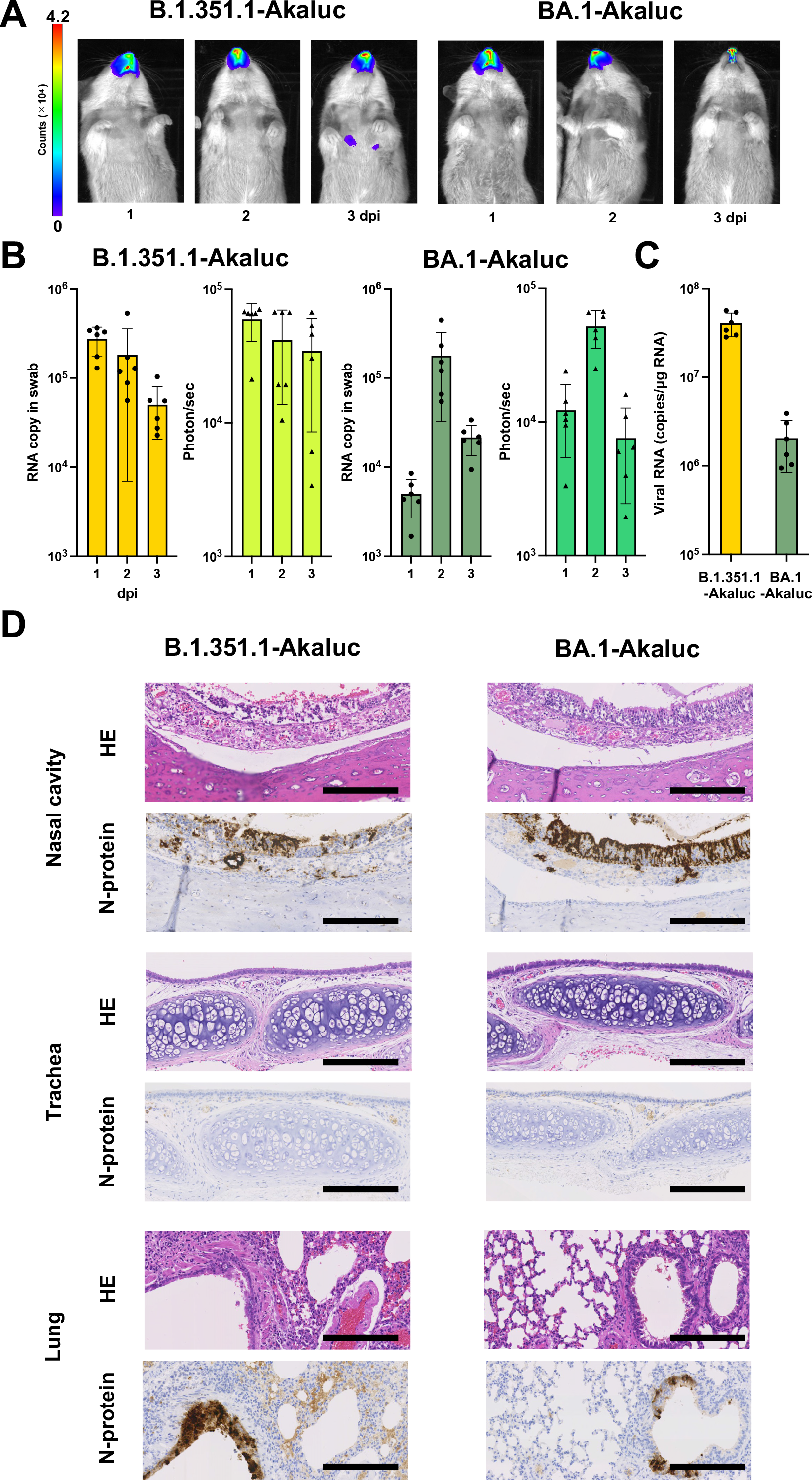
The virological features of SARS-CoV-2–Akaluc variants *in vivo*. Syrian hamsters (*n*=6 per group) were intranasally inoculated with B.1.351.1-Akaluc and BA.1-Akaluc. **(A)** Akaluc bioluminescence imaging of hamsters inoculated with the two viruses was performed daily through 3 dpi. Representative images are shown. **(B)** Viral RNA was quantified in oral swabs and the luminescence intensity of the nasal cavity of hamsters was measured daily through 3 dpi. **(C)** At 3 dpi, the lung hilum was harvested from hamsters and the viral RNA levels quantified. **(D)** IHC of the viral N protein (stained brown) was assessed in the nasal cavity, trachea, and lungs at 3 dpi. Representative figures are shown. Scale bars:100 µm. HE, hematoxylin and eosin

### Evaluation of *in vivo* efficacy against licensed SARS-CoV-2 antiviral and vaccine by AkaBLI

Several studies, including ours, showed that the Omicron subvariant XBB.1.5 has evolved to escape humoral immunity against the ancestral SARS-CoV-2 infection and/or vaccination.^30-32^ Thus, to examine whether AkaBLI can evaluate the immunoprophylactic capacity of antiviral or vaccine against SARS-CoV-2 variants, we generated recombinant, Akaluc-carrying SARS-CoV-2 encoding the XBB.1.5 S protein (XBB.1.5-Akaluc). We administered neutralizing monoclonal antibodies (AZD7442)^11^ currently used in humans to hamsters one day prior to inoculation with either B.1.1-Akaluc or XBB.1.5-Akaluc (*n*=6 per group) (**Figure 4A**). Unlike the hamsters who received the isotype control antibody, those who received AZD7442 prior to inoculation with B.1.1-Akaluc exhibited diminished Akaluc signals in the nasal cavity and no signals in the lungs (**Figure 4B**, upper panels), indicating AZD7442 provided protection from viral spread. In contrast, Akaluc signals in hamsters challenged with XBB.1.5-Akaluc did not differ depending on antibody treatment (**Figure 4B**, bottom panels), and viral RNA levels in the lungs were comparable between the two groups (**Figure 4C**).

**Figure 4.**
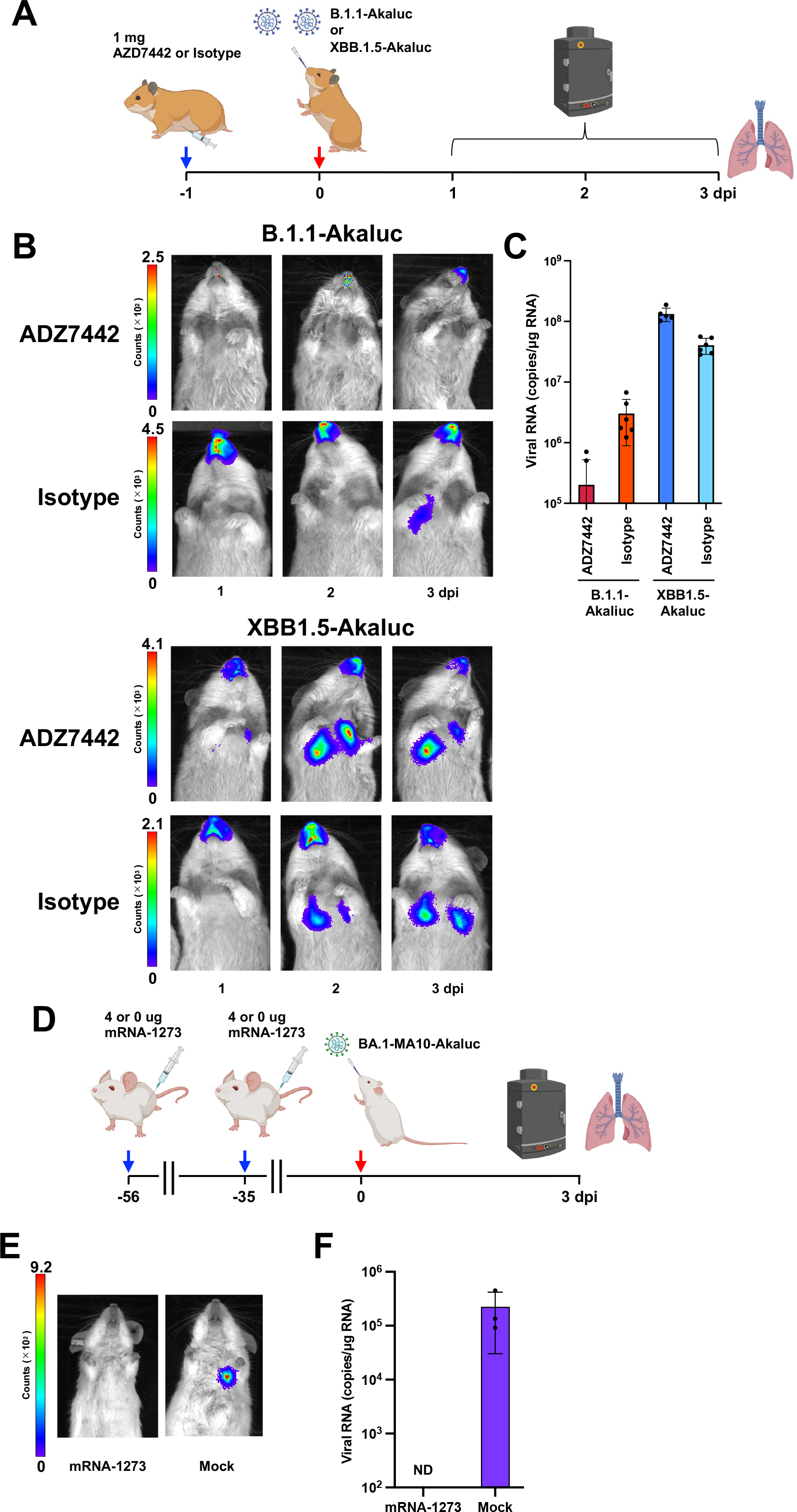
Evaluation of the immunoprophylactic ability of SARS-CoV-2 monoclonal antibodies and mRNA vaccine. **(A)** Schematic diagram of the experimental timeline for the evaluation of AZD7442 (Tixagevimab-Cilgavimab). One day before challenge, hamsters were immunized by AZD7442 (1 mg per dose) or isotype as control. Hamsters were inoculated with B.1.1-Akaluc (*n*=6), or XBB.1.5-Akaluc (*n*=6). One hamster in the XBB.1.5-Akaluc cohort unexpectedly died during the experiment. **(B)** Akaluc bioluminescence imaging was performed daily through 3 dpi. **(C)** The lung hilum was harvested at 3 dpi and subjected to viral RNA quantification. **(D)** Schematic diagram of experimental timeline for the evaluation of the mRNA vaccine BNT162b2. BALB/c mice (*n* =6) were vaccinated (4 μg per dose) twice at the indicated timepoints, and the other BALB/c mice (*n* =5) were not vaccinated (were received PBS as a control). Fifty-six weeks after the first vaccination, mice were inoculated with MA10-Akaluc (carrying the S protein of BA.1). Neutralizing antibody titers are shown in **Figure S2F**. Bioluminescence imaging was performed at 3 dpi. **(E)** Representative images from infected mice (immunized and non-immunized) are shown. (**F**) The lung hilum was collected at 3 dpi and subjected to viral RNA quantification.

Finally, to examine the utility of AkaBLI for evaluating the immunoprophylactic effects of mRNA vaccination, we used mouse-adapted SARS-CoV-2 (MA10)^33^ to generate recombinant, Akaluc-carrying SARS-CoV-2 encoding BA.1 S (BA.1-MA10-Akaluc). Mice were immunized twice with the licensed mRNA vaccine BNT162b2^12^and efficiently produced neutralizing antibodies against SARS-CoV-2 (**Figure S2F**). Five weeks after the second vaccination, control and immunized mice were challenged with BA.1-MA10-Akaluc (**Figure 4D**). At 3 days post-challenge, Akaluc signal (**Figure 4E**) and viral RNA (**Figure 4F**) were detected in the lungs of control, but not immunized, mice. Taken together, these data indicate that AkaBLI can evaluate the effects of immunoprophylaxis and immunotherapy against SARS-CoV-2 variants.

## Discussion

The insertion of large foreign genes, including luciferase, into viral genomes can be challenging. In previous studies, including ours, foreign genes have been accommodated into the loci of SARS-CoV-2 that encode accessory viral proteins.^13,26,28^ Thus, we attempted to replace viral ORFs 6-8 with the Akaluc gene but were unable to recover Akaluc-carrying viruses after transfecting cells with this modified genome (**Figure 1B**). Since the GC content of SARS-CoV-2 RNA is as low as 38% and that of the Akaluc gene is greater, we hypothesized that the GC ratio could affect the stability of foreign genes in the viral genome. We engineered a mutant Akaluc carrying synonymous substitutions to lower the gene’s GC content (**Figure S1**). The recombinant SARS-CoV-2 carrying this modified Akaluc exhibited robust replication in cell culture with infection kinetics similar to the parental B.1.1 genome (**Figure 1D**). These data suggest that the GC content of foreign genes should match that of the viral genome since GC content plays a critical role in viral propagation. Further investigations are needed to reveal the complete picture of the effect of GC content on the viral life cycle.

Several groups have reported that visualization of the *in vivo* dynamics of SARS-CoV-2 is possible with NanoLuc BLI in hACE2 transgenic mice^25,26^ and mice engrafted with human fetal lung xenografts in the skin.^27^ However, in the present study, NanoLuc signals could only be detected by IVIS in cell culture (**Figure S2C**) and not in hamsters infected with recombinant SARS-CoV-2 carrying NanoLuc (**Figure 1E**). This is because overexpression of hACE2^34^, the primary receptor of SARS-CoV-2, in transgenic mice enables robust viral replication not only in the respiratory tissues but also in nontarget organs. On the other hand, AkaBLI enabled spatio-temporal observation of recombinant SARS-CoV-2 infection dynamics in hamsters without exogenous overexpression of hACE2. These results indicate that AkaBLI is ideal for *in vivo* imaging of SARS-CoV-2 infection.

The insertion of the Akaluc gene did impair SARS-CoV-2 infection in hamsters (**Figure 2A** and **S2B**). In this study, we replaced ORFs 6-8 with the Akaluc gene. These accessory genes are dispensable for viral replication but are reported to be involved in immune response and pathogenicity.^35^ Accordingly, the lack of these genes in our B.1.1-Akaluc may have resulted in the attenuated pathogenicity we observed in hamsters. Notably, in the respiratory organs of the hamsters, Akaluc signals were correlated with the levels of viral RNA in the oral swabs and reflected viral dynamics (**Figure 2B** and **2C**).

Numerous studies, including ours, have shown that the tissue tropism and replication properties of SARS-CoV-2 have changed during circulation among humans.^10,17-24^ In **Figure 3**, the recombinant viruses carrying the S of the variants B.1.351.1 and BA.1 displayed different replication dynamics in hamsters and this was reflected by AkaBLI. Staining of viral N protein by IHC was correlated with the strength of the Akaluc signals we observed (**Figure 2D** and **3D**). Collectively, these data indicate that the AkaBLI can also be utilized to assess viral tissue tropism *in vivo*.

Most humans have experienced COVID-19 infection and/or been vaccinated, and SARS-CoV-2 variants are continuously emerging (reviewed in ^36^). Thus, our ability to protect against new variants is crucial for controlling the spread of infection. To assess whether AkaBLI can be used to evaluate antivirals, we assessed the effects of the neutralizing monoclonal antibodies AZD7442 or an mRNA vaccine on infection. Consistent with a previous report,^37^ AZD7442 protected against B.1.1 but XBB.1.5 *in vivo* (**Figure 4B** and **4C**). In addition, to evaluate immunoprophylaxis in mice, we generated a recombinant Akaluc-carrying, mouse-adapted SARS-CoV-2^33^. AkaBLI was able to visualize infection with this virus in mice. As shown in **Figures 4D** and **4E**, the mRNA vaccine fully protected mice against mouse-adapted SARS-CoV-2 infection. These findings suggest that AkaBLI can be used to investigate the *in vivo* efficacy of antivirals.

In summary, we constructed recombinant SARS-CoV-2 carrying the Akaluc gene and found that using AkaBLI we could monitor viral dynamics *in vivo*, evaluate differences in tissue tropism, and assess the efficacy of antivirals. Our findings will contribute to further studies in the engineering of recombinant viruses, including those of other RNA as well as DNA viruses, that can be used for *in vivo* studies. The development of novel biological assays such as AkaBLI is necessary to improve our understanding of the molecular mechanisms underlying virus replication and pathogenesis as well as to preclinically evaluate new antiviral agents.

## STARMethods

### Ethics statement

All experiments with hamsters and mice were performed in accordance with the Science Council of Japan’s Guidelines for the Proper Conduct of Animal Experiments. The protocols were approved by the Institutional Animal Care and Use Committee of National University Corporation Hokkaido University (approval number 20-0123).

### Plasmids

To generate recombinant SARS-CoV-2, 9 plasmids were used to amplify SARS-CoV-2 cDNA fragments (F1 to F9-10) by PrimeSTAR GXL DNA polymerase (Takara Bio) with the previously reported primer sets.^13^ The amplified DNA fragments were used for circular polymerase extension reaction (CPER; see below). To generate viruses carrying the NanoLuc luciferase gene, nucleotides 27,433–27,675 in the ORF7 region of pcDNA.3.1-CoV-2-G9-10 were replaced with the NanoLuc gene as previously described,^28^ and the resulting plasmids were designated as pcDNA.3.1-CoV-2-G9-10-NanoLuc. To generate viruses carrying the Akaluc luciferase gene (1,653 nt length)^7^, the nucleotide sequences of ORFs 6-8 in pcDNA.3.1-CoV-2-G9-10 were replaced with the Akaluc gene, and the resulting plasmids were designated as pcDNA.3.1-CoV-2-G9-10-Akaluc. Codon-optimized Akaluc was synthesized and purchased (Eurofins Scientific). SARS-CoV-2 B.1.1 Spike carrying a D614G mutation^38^ or Spike with B.1.351, BA.1^20^, or XBB.1.5^32^ sequence were inserted into a pCSII or pMW vectors, and were designated as pCSII-CoV-2-G8-D614G, pMW-CoV-2-G8-B.1.351, pMW-CoV-2-G8-BA.1, pMW118-CoV-2-G8-XBB.1.5. To generate mouse-adapted SARS-CoV-2 (MA10), which has seven mutations (C9438U, A11847G, A12159G, C23039A, U27221C, Q498Y, P499T) to facilitate binding to murine ACE2^33^, these mutations were incorporated into F3 and F4 and inserted into a pMW118 vector (pMW118-SARS-CoV-2-G3-MA10, -G4-MA10). Sequences of all the plasmids used in this study were confirmed by sequencing with SeqStudio Genetic Analyzer (Thermo Fisher Scientific) and outsourced service (Fasmac).

### Cell lines

All mammalian cell lines were cultured at 37°C under the conditions of a humidified atmosphere and 5% CO_2_. HEK293 (human embryonic kidney-derived 293)-3P6C33 cells (HEK293 cells stably expressing human ACE2 and TMPRSS2)^15^ were maintained in high-glucose Dulbecco’s Modified Eagle Medium (DMEM) (Nacalai Tesque) supplemented with 100 U/mL penicillin, 100 μg/mL streptomycin (PS) (Nacalai Tesque), and 10% fetal bovine serum. African green monkey kidney-derived VeroE6/TMPRSS2 cells (VeroE6 cells stably expressing human TMPRSS2)^14^ were maintained in low-glucose DMEM (FUJIFILM Wako Chemicals) supplemented with 1mg/mL G418 (Nacalai Tesque) and 10% fetal bovine serum.

### Preparation of viruses by CPER

Recombinant SARS-CoV-2 was generated by CPER as previously described.^13^ In brief, the 9 fragments of SARS-CoV-2 and UTR linker for SARS-CoV-2 described above were prepared by PCR using PrimeSTAR GXL DNA polymerase (Takara Bio). To prepare recombinant SARS-CoV-2 expressing the genes of MA10, B.1.351.1, BA.1, XBB.1.5, NanoLuc, or Akaluc mutations, the respective F3, F4, F8, or F9-10 plasmids were used for CPER. To generate recombinant SARS-CoV-2, the CPER products (25μl) were transfected into HEK293-3P6C33 cells with Opti-MEM (Thermo Fisher Scientific) and TransIT-LT1 (Mirus Bio) according to the manufacturer’s protocol. At 6 hours post-transfection, the culture medium was replaced with DMEM containing 2% FBS, 1% PS, and doxycycline (1 μg /mL) (InvivoGen). All viruses were stored at -80°C until use.

### Northern blotting

Total RNAs were extracted from cells infected with the wild-type SARS-CoV-2 (B.1.1) or SARS-CoV-2 carrying Akaluc and were subjected to Northern blot analysis as previously described^13^. In brief, a digoxigenin (DIG)-labeled random-primed probe, corresponding to 28,999 to 29,573 of the SARS-CoV-2 genomes, was generated by using a DIG RNA Labeling kit (SP6/T7) (Sigma-Aldrich), and utilized to detect viral mRNAs. The RNAs were washed with the DIG luminescent detection kit (Sigma-Aldrich) and were visualized with CDP-Star Chemiluminescent Substrate (Sigma-Aldrich), according to the manufacturer’s protocols. Bands were detected by WSE-6100LuminoGraphI (Atto).

### Quantitative RT-PCR

For quantification of viral RNA copies, total RNA was extracted from cells using a PureLink RNA Mini Kit (Thermo Fisher Scientific), and then first-strand cDNA synthesis and qRT-PCR were performed using One Step PrimeScript III RT-PCR Kit (Takara Bio) and QuantStudio 5 Real-Time PCR System (Thermo Fisher Scientific), respectively, according to the manufacturer’s protocols. For quantification of viral RNA, primer sets detecting the region encoding part of the viral nucleocapsid phosphoprotein as reported in a previous study^15^ were used. Fluorescent signals were determined by QuantStudio 5 Real-Time PCR System.

### Titration and growth kinetics

The infectious titers in culture supernatants were determined by quantifying the 50% tissue culture infectious dose (TCID_50_).^39^ Cell culture supernatants were used to inoculate naive VeroE6/TMPRSS2 cells in 96-well plates after ten-fold serial dilution with DMEM containing 1mg/mL G418 (Nacalai Tesque) and 2% FBS, and the infectious titers were determined at 72 hours post-infection (hpi). For growth kinetics, SARS-CoV-2 was inoculated into VeroE6/TMPRSS2 cells in 6-well plates at a multiplicity of infection (MOI) of 0.01, the culture supernatants were replaced with new medium at 1 hpi and incubated for 48 h. The infectious titers ofthe culture supernatants collected at 12, 24, 36, and 48 hpi were then determined.

### Passage experiment

One day before infection, 300,000 HEK293-3P6C33 cells/well were seeded into a 6-well plate. SARS-CoV-2 carrying Akaluc was used to inoculate the HEK293-3P6C33 cells at an MOI of 1.0. At 1 hpi, the culture supernatants were replaced with new medium and then incubated for 48 h. Culture supernatants collected at 48 hpi were used as an inoculum for new HEK293-3P6C33 cells and also used for viral RNA extraction and sequence analysis (See **Figure S2A, Table S1**).

### Bioluminescence experiment

Luminescence was detected using AkaLumine-HCl (FUJIFILM Wako Chemicals), and luciferase activity *in vitro* was measured using an AB-2270 Luminescencer Octa (Atto) according to the manufacturer’s protocol. VeroE6/TMPRSS2 cells were seeded in a black 96-well plate (PerkinElmer) at a density of 15,000 cells per well. Serially diluted wild-type B.1.1 or recombinant B.1.1 carrying NanoLuc or the Akaluc gene were used to infect the cells. At 32 hpi, cells were washed once with PBS and imaged using an IVIS Imager (PerkinElmer) at 5 min after addition of substrate: fluorofurimazine (FFz) (3 g/ml: Promega) for NanoLuc and AkaLumine-HCl (4.3 g/ml) for Akaluc.

For *in vivo* experiments, infected hamsters or mice were injected with FFz (440 nmol/g) or AkaLumine-HCl (75 nmol/g) intraperitoneally according to the indicated protocol from the previous studies^7,29^ and the luminescence images were monitored by IVIS at 1, 2, and 3 dpi. All images were analyzed using Living Image^®^ Ver4.7.3 (PerkinElmer).

### Animal experiments

Animal experiments were performed as previously described.^17-24^ Syrian hamsters (male, 4 weeks old) were purchased from Japan SLC Inc. BALB/c mice (male, 10 weeks old) were purchased from CLEA Japan Inc. For the virus infection experiments, animals were anesthetized by intramuscular injection with a mixture of 0.15 mg/kg medetomidine hydrochloride (Domitor, Nippon Zenyaku), 2.0 mg/kg midazolam (Dormicum, Fujifilm WAKO Chemical), and 2.5 mg/kg butorphanol (Vetorphale, Meiji Seika Pharma) or 0.15 mg/kg medetomidine hydrochloride, 4.0 mg/kg alphaxaone (Alfaxan, Jurox), and 2.5 mg/kg butorphanol. Animals were intranasally inoculated under anesthesia with the recombinant viruses (10,000 TCID_50_ in 100μl for hamsters or 1,000 TCID_50_ in 50μl for mice) or saline (100 µl). Oral swabs were collected at the indicated timepoints. Body weight was recorded daily through 7 dpi. Lung tissues were anatomically collected at 3 dpi. The viral RNA load in the oral swabs and respiratory tissues was determined by quantitative RT–PCR. These tissues were also used for IHC and histopathological analyses (see below). For evaluation of *in vivo* efficacy against licensed SARS-CoV-2 antiviral and vaccine, hamsters were intramuscularly injected with 1 mg per dose of AZD7442 (Tixagevimab-Cilgavimab; AstraZeneca) one day before viral infection. Mice were received twice with 4 μg of the mRNA vaccine BNT162b2 (Pfizer/BioNTech) 7 and 5 weeks before virus challenge.

### IHC

IHC was performed as previously described^17-24^ using an Autostainer Link 48 (Dako). The deparaffinized sections were exposed to EnVision FLEX target retrieval solution high pH (Agilent) for 20 minutes at 97°C for activation, and a mouse anti-SARS-CoV-2 N monoclonal antibody (clone 1035111, R&D Systems, dilution 1:400) was used as a primary antibody. The sections were sensitized using EnVision FLEX for 15 min and visualized by peroxidase-based enzymatic reaction with 3,3′-diaminobenzidine tetrahydrochloride (Dako) as substrate for 5 minutes. Images were incorporated as virtual slides by NDP.scan software v3.2.4 (Hamamatsu Photonics).

### Statistical analysis

All assays were performed in triplicate and independently repeated at least two times. Results were expressed as the means±standard deviations or standard errors. Statistical significance was determined by the two-tailed Student’s *t*-test performed by GraphPad Prism (Software ver. 10.0.1). Significantly different values (*p*<0.05) are indicated by an asterisk.

## Author contributions

T.T., Shiho T., and T.F. designed the research; T.T., H.I., Shiho T., L.W., R.S., Shuhei T., A.K., and Yu. M. performed the research; T.T., S.S., Kotaro. S., Kei S., Yo. M., Shinya T., S.I., and T.F. analyzed the data; and T.T., H.I, and T.F. wrote the paper.

Japan Agency for Medical Research and Development

## Acknowledgements

We thank H. Kubo, M. Tetsuka, and S. Shimamura for their secretarial work and H. Maruyama, M. Hanazaki, T. Matsuoka, A. Imajoh, K. Oyama, and H. Murota for their technical assistance. This work was supported in part by AMED SCARDA Hokkaido University Institute for Vaccine Research and Development (HU-IVReD) (JP23fa627005h0001 to Takasuke Fukuhara); AMED SCARDA (JP223fa827001h0001 to Takasuke Fukuhara), AMED Research Program on Emerging and Re-emerging Infectious Diseases (JP21fk0108617h0001, JP21fk0108493h0001, JP22fk0108511h0001, and JP22fk0108516h0001 to Takasuke Fukuhara); AMED Project for Advanced Drug Discovery and Development (JP21nf0101627 to Takasuke Fukuhara); AMED CREST (JP22gm1610008h0001 to Takasuke Fukuhara); JSPS KAKENHI Grant-in-Aid for Scientific Research B (21H02736 to Takasuke Fukuhara); Takeda Science Foundation (to Takasuke Fukuhara); and Hokkaido University Support Program for Frontier Research (to Takasuke Fukuhara).

## Supplemental information

**Supplementary Figure 1.**
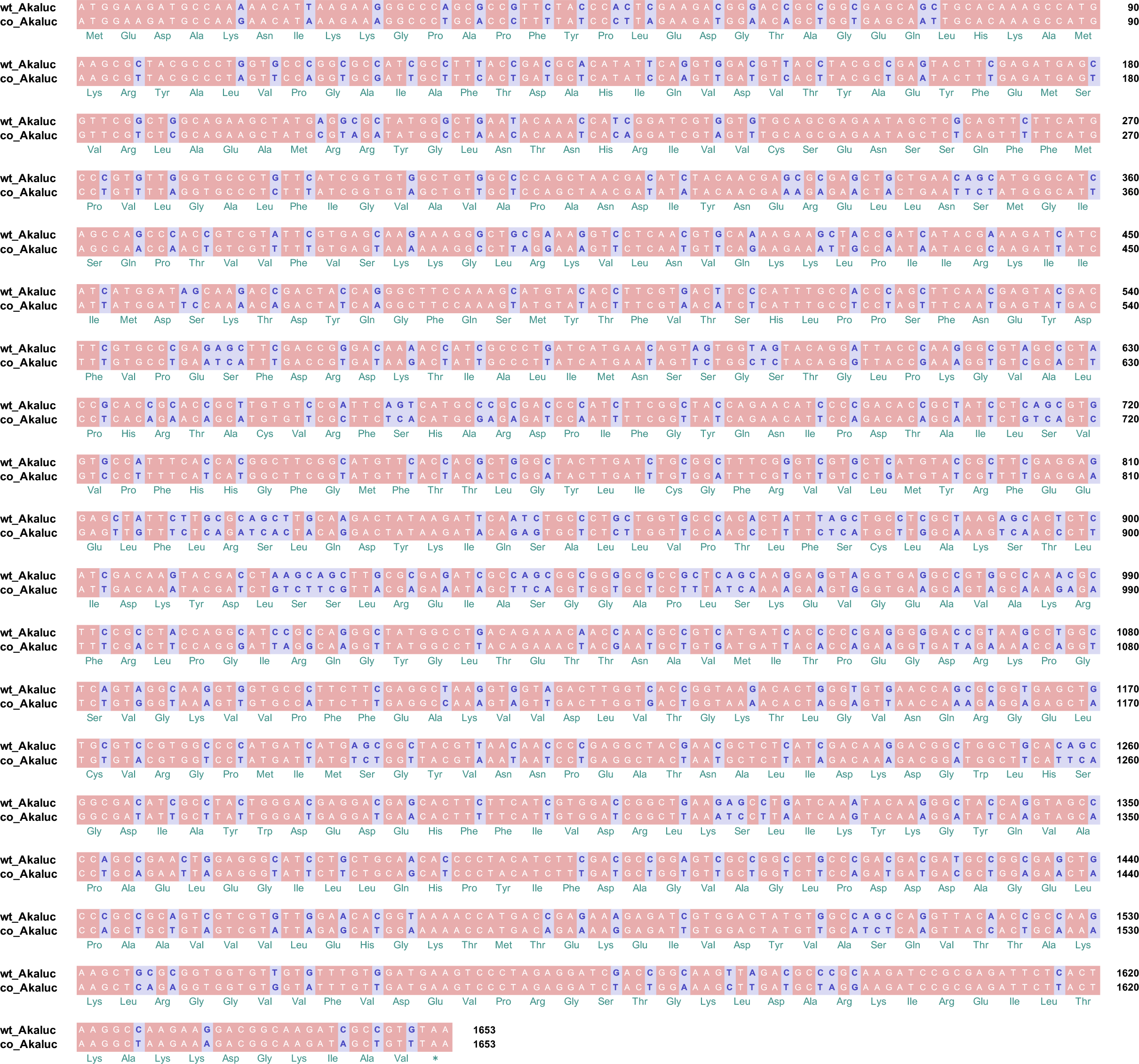
Nucleotide alignment of the wild-type and codon-optimized Akaluc luciferase genes. wt_Akaluc: wild-type Akaluc gene (GenBank accession number: LC320664.1); co_Akaluc: codon-optimized Akaluc gene.

**Supplementary Figure 2.**
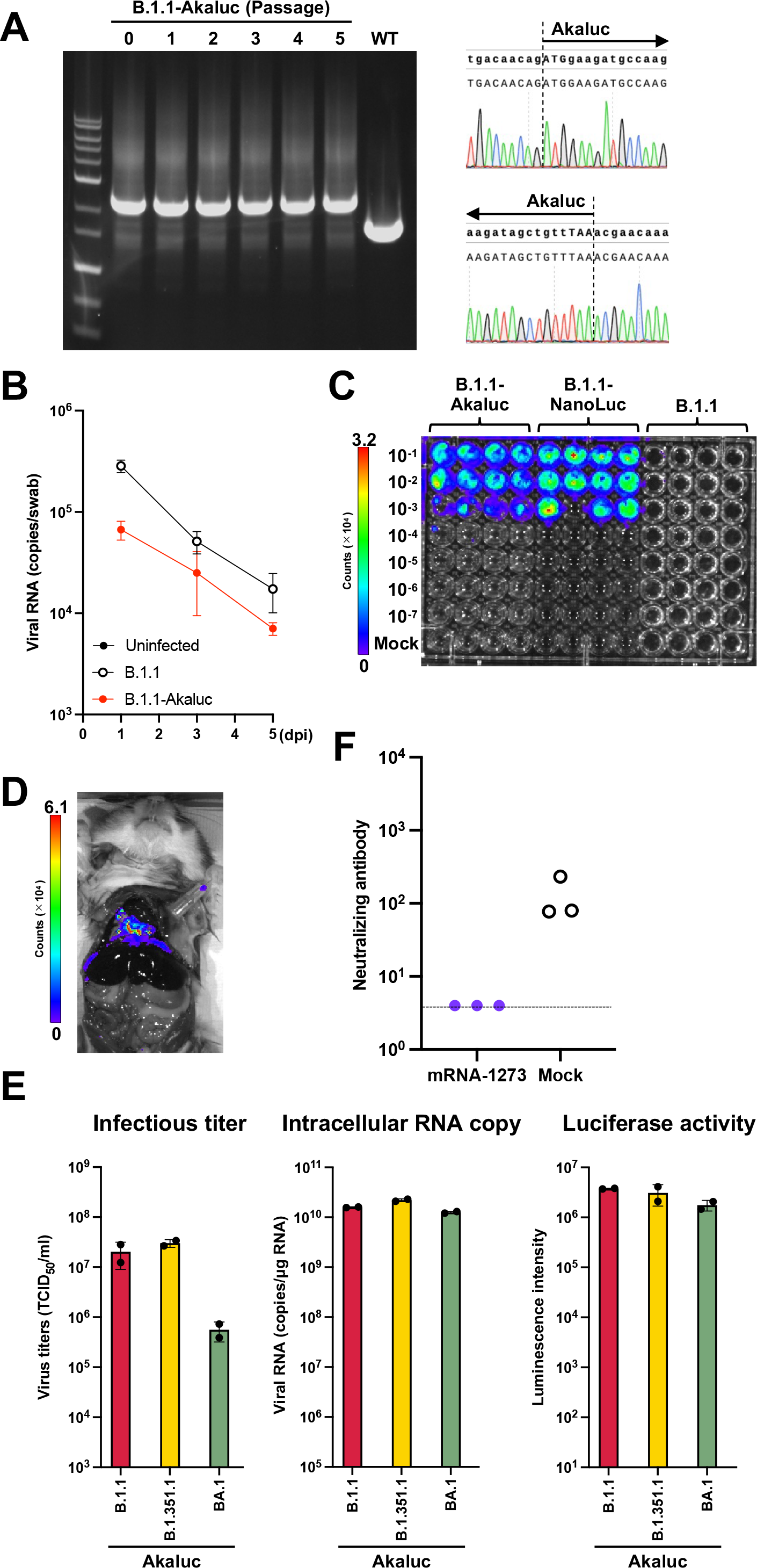
Characterization of SARS-CoV-2 reporters. **(A)** Stability of the Akaluc luciferase gene in SARS-CoV-2 Akaluc. HEK293-3P6C33 cells were infected (MOI=1.0) with B.1.1-Akaluc (P0). Culture supernatants were harvested at 48 hpi (P1) and used to inoculate naive cells. This procedure was repeated through P5. Viral RNA was extracted from the P5 supernatants and the Akaluc gene was amplified by PCR. These fragments were subjected to electrophoresis (left) and sequence analysis (right). Sequence results for the Akaluc gene junction are shown. **(B)** Viral RNA levels in oral swabs from hamsters infected with SARS-CoV-2 B.1.1 or B.1.1-Akaluc (n=6 per infection group). **(C)** Serial dilutions B.1.1, B.1.1-Akaluc or B.1.1-NanoLuc were used to inoculate VeroE6/TMPRSS2 cells. Cultures were imaged by BLI at 32 hpi. **(D)** Representative bioluminescence imaging of the exposed pleural cavity of a hamster infected with B.1.1-NanoLuc. **(E)** *In vitro* properties of SARS-CoV-2–Akaluc recombinant viruses carrying the S protein of different SARS-CoV-2 variants. VeroE6/TMPRSS2 cells were infected with the viruses indicated (MOI=0.01). Infectious titers of the culture supernatants (left panel), intracellular viral RNA copy numbers (middle panel), and luciferase activities (right panel) were determined at 36 hpi. **(F)** Neutralizing antibodies against SARS-CoV-2 B.1.1 of the immunized and non-immunized mice in the serum.

**Supplementary Table 1. Mutations in B**.**1**.**1-AkaLuc after serial passaging**.

